# Novel microRNA-455-3p mouse models to study Alzheimer’s disease pathogenesis

**DOI:** 10.1101/2021.09.23.461513

**Authors:** Subodh Kumar, Hallie Mortan, Neha Sawant, Erika Orlov, Lloyd Bunquin, Jangampalli Adi Pradeepkiran, P. Hemachandra Reddy

## Abstract

MicroRNA-455-3p is one of the highly conserved miRNAs involved in several human diseases but newly explored by our lab in Alzheimer’s disease (AD). Our past studies unveiled the biomarker and therapeutic potentials of miR-455-3p in AD. Our *in vitro* study exhibited the protective role of miR-455-3p against AD toxicities in reducing full-length APP and amyloid-β (Aβ) protein levels, and also reducing defective mitochondrial biogenesis, impaired mitochondrial dynamics and synaptic deficiencies. Next, we sought to determine the essential roles of miR-455-3p in AD using mouse models. Therefore, for the first time we generated both transgenic (TG) and knockout (KO) mouse models of miR-455-3p. We determined the positive and negative effects of miR-455-3p on mice cognitive function, mitochondrial biogenesis, mitochondrial dynamics, mitochondrial number & length, dendritic spine density, synapse numbers and synaptic activity in 12-month-old miR-455-3p TG and KO mice. MiR-455-3p TG mice lived 5 months longer than wild-type (WT) mice, whereas KO mice lived 4 months shorter than their WT counter parts. Morris water maze test showed improved cognitive behavior, spatial learning and memory in miR-455-3p TG mice relative to age-matched WT mice and miR-455-3p KO mice. Further, mitochondrial biogenesis, dynamics and synaptic activities were enhanced in miR-455-3p TG mice, while these were reduced in KO mice. Overall, miR-455-3p TG mice displayed protective effects and miR-455-3p KO mice exhibited deleterious effects in relation to AD pathogenesis. Both mouse models could be ideal research tools to understand the molecular mechanism of miR-455-3p in AD and other human diseases.

## Introduction

Alzheimer’s disease (AD) is the most common dementia that progressively decline the memory and other mental functions in elderly individuals [1]. Currently, over 50 million people worldwide live with AD-related dementia, and this number is expected to increase to 152 million by 2050 (World Alzheimer Report 2020). AD is manifested by dementia in aged individuals [1]. AD-related dementia has huge economic consequences, with the total medical costs of dementia worldwide is estimated at $818 billion (World Alzheimer Report 2020). In addition to dementia, AD is associated with the loss of synapses, synaptic dysfunction, microRNA (miRNA) deregulation, mitochondrial structural and functional abnormalities, inflammatory responses, extracellular neuritic plaques, and intracellular neurofibrillary tangles (NFTs) [1–11]. Despite tremendous progress that has been made in better understanding the AD pathogenesis, there are still no detectable markers, drugs, and/or agents that can prevent AD or slow its progression. Search for a novel, non-invasive, early detectable biomarkers that are involved in AD and Alzheimer’s disease related disorders (ADRD) still ongoing. Several miRNAs have been reported that are involved in AD-related pathways and regulate AD pathogenesis via modulation of AD genes [6, 12–18]. However, most of the studies are limited to *in vitro* cell culture treatment only [13–20]. MiRNA based intervention is the best way to study and develop therapeutics in AD. To understand the physiological functions of miRNA, it is important to generate both TG and KO mouse model, particularly for miRNAs associated with APP processing and amyloid beta (Aβ).

MiR-455 is a member of a broadly conserved miRNA family expressed in most of phyla, including Mammalia and Primates. MiR-455 mostly implicated in human cancers and chondrogenic differentiations [21]. Recent reports unveiled the role of miR-455-3p in mitochondrial biogenesis through the upregulation of the PGC1α gene via regulation of novel HIF1an-AMPK-PGC1α, a signaling network [22]. For the first time, we investigated the connection of miR-455-3p in AD. Our lab was the first to investigate the relevance of miR-455-3p in AD [21, 24–26]. Our global microarray analysis found higher expression of miR-455-3p in the serum samples from patients with AD relative to healthy controls [23]. Further, findings on AD postmortem brains, AD-Fibroblasts, AD B-lymphocytes (all from late onset AD; LOAD), AD cerebrospinal fluids, AD cell lines, and APP transgenic (Tg2576 strain) mice exhibited higher expression levels of miR-455-3p in AD cases compared to the samples from healthy controls [21, 23–26].

MiR-455-3p target several genes, most of them validated in cancers [21]. In relation to AD, APP was the most authenticated and one of the top predicted targets of miR-455-3p [25]. Overexpression of miR-455-3p construct reduces the expression of mutant APP cDNA in mouse neuroblastoma cells. The levels of full-length APP, C-terminal fragments of APP (C99 and C83) and the levels of Aβ1-40 and Aβ1-42 were also significantly decreased in cells by miR-455-3p. Further, overexpression of miR-455-3p reduces the toxic effects of Aβ on mitochondrial biogenesis, mitochondrial dynamics, synaptic activities, cell viability and apoptosis [25].

Based on the *in vitro* findings, it is well established that high levels of miR-455-3p reduces Aβ pathology, enhances mitochondrial biogenesis, mitochondrial function, and synaptic activity, whereas our studies of reduced the endogenous miR-455-3p, showed increased Aβ, reduced mitochondrial biogenesis and reduced mitochondrial function. However, to confirm our *in vitro* findings, and understand the physiological functions of miR-455-3p, creation of overexpressed (TG) and depleted (KO) miR-455-3p mouse models is necessity. Mouse models are the best tools to study the molecular mechanism and impact of miRNAs in the disease process. <u>Therefore, in the current study, using pronuclear injection and CRISPR technologies, for</u> <u>the first time, we generated both transgenic and knockout mouse models of miR-455-3p and</u> <u>characterized for lifespan extension, cognitive behavior, mitochondrial and synaptic activities.</u>

## Materials and methods

### Animals

The animal study was approved by Texas Tech University Health Sciences Center - Institutional animal care and use committee (TTUHSC-IACUC). The wild type (WT) mice (C57BL6/J) aged 12 months were purchased from The Jackson Laboratory (Bar Harbor, ME). The microRNA-455-3p Transgenic (miR-455-3p TG; TG) and microRNA-455-3p Knockout (miR-455-3p KO; KO) mouse models were generated in collaboration with Cyagen Biosciences, Santa Clara, CA, USA. The twelve months old mice were divided into three groups-Group 1: WT mice, Group 2: TG, and Group 3: KO heterozygous (KO+/−), we call these animals KO, from here on in the manuscript. All animals were housed under air-conditioned rooms at a constant temperature of 22°C with a 12 h light/dark cycle and given access to water and food ad libitum.

### Generation of miR-455-3p transgenic mouse model

MiR-455-3p overexpressing TG mouse model was generated on a C57BL6 mice background, using a pronuclear plasmid injection of miR-455-3p expression vector (pRP[Exp]-U6>hsa-miR-455-3p-CAG-EGFP) as shown in supplementary information Figure 1 (SI Fig. 1) in collaboration with Cyagen Biosciences (Supplementary materials). Before, the generation of TG mice, the biological properties of miR-455-3p construct were tested *in vitro* using mouse neuroblastoma cells [25]. To generate TG model, we tagged miR-455-3p construct with the green fluorescence protein (GFP), in order to detect miR-455-3p transfection and the localization of GFP within cells [25]. The resulting TG mice founders did not show any kind of physical abnormality and lethality related to miR-455-3p transgene at any stage of life and TG animals viable and fertile.

**Figure 1.**
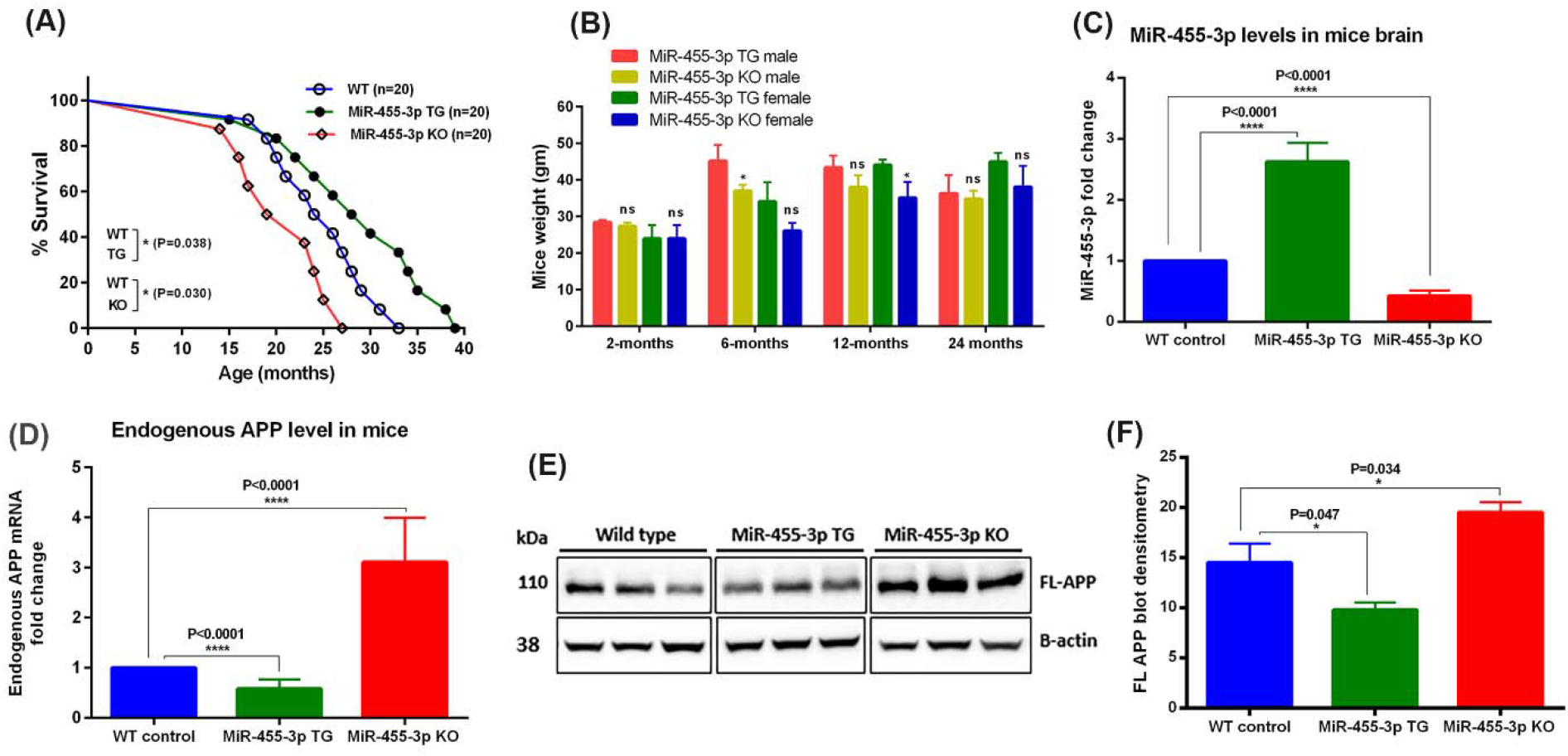
**A-** Kaplan-Meier survival curve of WT, miR-455-3p TG and miR-455-3p KO mice. The survival percentage was significantly increased (P=0.038) in TG mice (n=20) relative to WT mice (n=20). In KO mice (n=20) survival percent was significantly decreased (P=0.030) relative to WT mice. **B-** Mice weight comparisons analysis in miR-455-3p TG and miR-455-3p KO mice. Mice weights (gm) were calculated for - 2 months, 6 months, 12 months and 24 months periods in both males and females. The significant differences (*P<0.05) in the mice weight was seen in 6-months KO male relative to 6 months TG male and 12 months old KO female relative to 12 months TG female. **C-** qRT-PCR analysis of miR-455-3p expression in TG (n=15) and KO (n=15) mice relative to age and sex matched WT mice (n=15). MiR-455-3p fold change was significantly higher (P<0.0001) in TG mice relative to WT mice. In KO mice miR-455-3p fold change was significantly lowered (P<0.0001) relative to WT mice. **D-** qRT-PCR analysis of mouse endogenous APP mRNA expression in TG (n=15) and KO (n=15) mice relative to age and sex matched WT mice (n=15). APP mRNA fold change was significantly reduced (P<0.0001) in TG mice relative to WT mice. In KO mice APP mRNA fold change was increased significantly (P<0.0001) relative to WT mice. **E-** Immunoblot analysis for mouse full-length endogenous APP in WT (n=5), TG (n=5) and KO (n=5) mice brain. Immunoblots showing the levels of FL-APP (110 kDa) and B-actin (38 kDa) in three representative mice in all three groups of mice. **F-** Densitometry analysis for mouse FL-APP in WT, TG and KO mice. The level of APP protein was significantly lower (P=0.047) in TG mice relative to WT mice. In KO mice protein level was significantly increased (P=0.034) relative to WT mice.

### Generation of miR-455-3p knockout mouse model

MiR-455-3p KO (+/−) mouse model for miR-455-3p suppression was generated using the CRISPR/Cas9 technology to knockout the endogenous miR-455-3p (Supplementary materials). Two pairs of gRNA constructs targeting the 3’ of mouse miR-455 and Cas9 mRNA were co-injected into fertilized mouse eggs to generate targeted knockout offspring (SI Fig. 2). F0 founder animals were identified by PCR, followed by sequence analysis, which were bred to wild type mice to test germline transmission and F1 animal generation. The deletion of 41 bp sequence of miR-455-3p was confirmed by sequencing analysis (SI Fig. 3). The resulting miR-455-3p KO positive founders were characterized by KO specific genotyping strategy (SI Fig. 2). The miR-455-3p TG and miR-455-3p KO confirmed pups were bred to obtain more pups (SI Fig. 4). The miR-455-3p KO pups did not show any kind of physical abnormality and lethality related to miR-455-3p deletion at any stage of life.

### Analysis of lifespan extension

To determine the impact of miR-455-3p on mice survival, we assigned the 20 animals in each groups WT, TG and KO mice for their lifespan analysis. Mice were monitored, daily basis for health checks and other their phenotypic changes, if any. The veterinary staff routinely monitored the health disparities in these mice. We also assessed the mice weight over the period at 2 months, 6 months, 12 months and 24 months of age.

### Mice cognitive behavior analysis-Morris Water Maze test

To determine the impact of miR-455-3p on spatial long-term learning and memory, we performed MWM test in 12-months old WT (n=15), TG (n=15) and KO (n=15) mice. The MWM test was performed as per our published protocol [27,28]. Details of the tests are provided in supplementary information.

### Assessment of miR-455-3p expression

Total RNAs were extracted from cortices by TRIzol RNA isolation reagent (Invitrogen, USA) as per manufacturer instructions. Quantification of miR-455-3p involved three steps: (i) Polyadenylation, (ii) cDNA synthesis and (iii) qRT-PCR. Total RNA was polyadenylated with a miRNA First-Strand cDNA synthesis kit (Agilent Technologies Inc., CA, USA), following manufacturer’s instructions. Primers used in the current study were synthesized commercially (Integrated DNA Technologies, Inc., City, Iowa, USA) (SI Table 1). The qRT-PCR analysis was performed as per our published protocol [23–26].

### Messenger RNA levels of mitochondrial biogenesis/dynamics and synaptic genes

Quantification of mRNA levels of mitochondrial biogenesis, dynamics and synaptic genes was carried out using real-time qRT-PCR using methods [25]. The oligonucleotide primers were designed with primer express software (Applied Biosystems) for the housekeeping genes β-actin, full-length mouse endogenous APP; mitochondrial biogenesis (PGC1α, Nrf1, Nrf2, TFAM: mitochondrial dynamics (Drp1, Fis1, Opa1, Mfn1, Mfn2) and synaptic & dendritic (SNAP25, PSD95 and MAP2) genes. The primer sequences and amplicon sizes are listed in SI Table 1.

### Immunoblot analysis of mitochondrial biogenesis, dynamics and synaptic proteins

Immunoblot analysis was performed, using protein lysates prepared from the cortices of all three groups of WT, TG and KO mice. We performed the immunoblot analysis for the mouse endogenous APP, mitochondrial biogenesis proteins (PGC1α, Nrf1, Nrf2, TFAM), mitochondrial dynamics proteins (Drp1 & Fis1- fission and Opa1, Mfn1 & Mfn2- fusion) and synaptic and dendritic proteins (SNAP25, PSD95 and MAP2). Details of the antibody dilutions are given in SI Table 2. Briefly, 40 μg of protein lysates were resolved on a 4-12% Nu-PAGE gel (Invitrogen) and transferred to PVDF membrane. After overnight incubation with the primary antibodies, the membranes were incubated for 2 h with an appropriate secondary antibodies. Proteins were detected with chemiluminescence reagents (Pierce Biotechnology, Rockford, IL, USA), and band intensities were quantified using NIH-image-J software [25].

### Immunostaining analysis

To determine the immunoreactivity and intensity of the cell type specific markers, mitochondrial biogenesis, dynamics and synaptic protein, we performed immunofluorescence analysis using brain sections of WT, TG and KO mice. The brains were sectioned coronally with a thickness of 10 μm size using the cryostats. Sections were washed with PBS and fixed in freshly prepared 4% paraformaldehyde for 10 min. After permeabilization, and blocking steps sections were incubated with primary antibodies (1:200 dilution) (SI Table 2). After secondary antibody incubation, photographs were taken with a multiphoton laser scanning microscope system (Olympus IX83 inverted microscope). To quantify the immunoreactivities of antibodies, 10-15 photographs were taken at 4X, 10X, 20X, 40X and 100X magnification. Statistical significance was assessed by the intensities of red, green or blue, using NIH ImageJ software [25].

### Golgi-cox staining and dendritic spine count

The morphology of neuronal dendrites and dendritic spines were observed in the brains of WT, TG and KO mice by Golgi-Cox staining, which was performed using the FD Rapid GolgiStain Kit (FD NeuroTechnologies, Columbia, MD, USA) as described earlier [27,28]. The dried brain sections were processed as per the manufacturer’s instructions. Briefly, dendrites within the CA1 sub region of the hippocampus were imaged using a 4X, 10X, 20X and 63X objective using EVOS microscope-AMG (thermofisher.com) and Olympus IX83. Dendritic spines were detected along CA1 secondary dendrites starting from their point of origin on the primary dendrite, and the counting was performed by an experimenter blinded of each group samples using Image J software [27,28].

### Transmission electron microscopy and mitochondrial number and length

To determine the effects of miR-455-3p on the mitochondrial number and size, we performed transmission electron microscopy in hippocampal and cortical sections of 12- month-old WT mice, TG and KO mice. Animals were perfused using saline standard perfusion method, and the brains were removed from the mice. The ventral part of the hippocampus layer-the CA1 region and cerebral cortex were isolated and cut into ∼1 mm 3 cubes. Tissues were fixed in a solution of 0.1M cacodylate buffer, 1.5% paraformaldehyde and 2.5% glutaraldehyde and then post-fixed with 1% osmium tetroxide and embedded in LX-112 resin. Ultrathin sections were cut, stained with uranyl acetate and lead citrate, and examined with a The Hitachi H-7650/Transmission Electron Microscope at 60 kV located at the College of Arts and Sciences Microscopy, Texas Tech University [25,27,28].

### Transmission electron microscopy and quantification of synapse numbers

Transmission electron microscopic images were used for the synapse quantification in cortex and hippocampal area of control WT, TG and KO mice brains. Images at 500 nm scale were selected for the quantification. Synapse numbers were counted with focus on the synapse organization (mitochondria, synaptic vesicles and synaptic cleft numbers) and average number of synapses in each slide were determined in WT, TG and KO brain.

### Statistical considerations

Statistical analyses were conducted, by using the student t-test for analyzing two groups of samples, and one-way comparative analysis of variance (ANOVA) was used for analyzing WT, TG and KO groups of mice data. Significant differences in three group of samples were calculated by Bonferroni’s multiple comparison tests. Statistical parameters were calculated, using Prism software, v6 (La Zolla, CA, USA). P<0.05 was considered statistically significant.

## Results

### Generation of miR-455-3p TG and miR-455-3p KO mouse models

The miR-455-3p TG and miR-455-3p KO mouse models were generated in collaboration with Cyagen Biosciences. The TG mice were generated by the pronuclear injection of miR-455-3p expression vector (pRP[Exp]-U6>hsa-miR-455-3p-CAG-EGFP) in the embryo of C57BL/6 mice. Over 5 founder lines of mice were generated, we continued to bred higher expresser line for further breeding and characterization. The TG positive pups were confirmed by transgene specific genotyping strategy.

MiR-455-3p KO mere were generated using the CRISPR/Cas-mediated genome engineering. The mouse miR-455 gene (miRBase: MI0004679; Ensembl: ENSMUSG00000070102) is located on the mouse chromosome 4. Briefly, 3’ P region of mouse miR-455 was selected as target site and two pairs of gRNA targeting vectors will be constructed and confirmed by sequencing (SI Fig. 2 and 3). Cas9 mRNA, gRNA generated by *in vitro* transcription were co-injected into fertilized eggs for KO mouse production. The resulted miR-455-3p KO positive F1 founders were genotyped and then sequence-analyzed to confirmed the depleted miR-455 locus of 41 total size (SI Fig. 3). The transgenes positive founders were characterized by genotype specific PCR for TG and KO mice (SI Fig. 4).

### Characterization of miR-455-3p TG and KO mouse models

The resulting miR-455-3p TG and KO mice were bred for further molecular characterizations. We studied the mice phenotypic behavior and overall survival of WT, TG and KO mice.

To determine the miR-455-3p expression-specific survival expectancy of miR-455-3p TG and KO mice, we studied the mice survival for lifetime. In each group, we assigned 20 mice for lifetime survival analysis. Based on Kaplan-Meier survival curve, the median survival was 25 months in WT mice, 29 months in TG mice and 21 months in KO mice. The TG mice lived 5 months longer than WT counter parts (P=0.038) and on the other hand, KO mice lived 4 months shorter than WT mice (P=0.030) (Fig. 1A). These observations, indicate that overexpression of miR-455-3p extend lifespan in mice whereas depletion of miR-455-3p reduces overall lifespan of mice.

The mice weight analysis showed the significant difference between 6 months old TG male vs KO male mice and 12 months old TG female vs KO female mice (Fig. 1B). The 6 months old KO male and 12 months KO female mice showed significantly reduced weight (*P<0.05) in relation to respective TG mice.

Next, we checked the expression of miR-455-3p and mouse endogenous full-length APP in both TG (n=15) and KO (n=15) mice brain. The qRT-PCR analysis showed the significant (P<0.0001) upregulation of miR-455-3p (Fig. 1C) and significant downregulation (P<0.0001) of mouse endogenous APP mRNA (Fig. 1D) in TG mice relative to WT mice. Opposite to that, miR-455-3p expression was significantly downregulated (P<0.0001) in KO (n=15) mice (Fig. 1C). As expected, miR-455-3p expression levels reduced to 50% in KO+/− mice. However, endogenous APP expression was significantly increased (P<0.0001) in KO mice relative to WT mice (Fig. 1D). These observations confirm the successful generation of miR-455-3p TG and KO mouse and regulation of APP gene by miR-455-3p at genetic level. Further, immunoblot analysis showed reduced levels of mouse endogenous APP proteins in miR-455-3p TG mice and increased levels in KO mice (Fig. 1E). Densitometry analysis showed significant reduction in the APP levels in TG mice relative to WT (P=0.047) and significant upregulation in KO mice relative to WT control mice (P=0.034) (Fig. 1F). All these observations confirmed the successful generation of miR-455-3p TG and KO mouse models.

### Impact of miR-455-3p on mice cognitive behavior, spatial learning and memory

To study the impact of miR-455-3p on mice cognitive behavior, we performed the Morris Water Maze (MWM) tests to access the spatial learning and memory in 12-month-old WT mice (n=15), TG mice (n=15) and KO mice (n=15). The significant differences were observed in MWM test, where miR-455-3p TG mice performed best relative to WT and KO mice (Fig. 2). We performed sixteen trials (5 training and 11 real) of MWM. As shown in Fig. 2A, the tracking plot (A) and heat map (B) showed the clear difference between MWM performance of WT, TG and KO mice. The average latency time to find the platform was significantly decreased in miR-455-3p TG mice compared to the control WT mice (P=0.037) and KO mice (P=0.003) (Fig. 2B). The total distance travelled by mice was also decreased in miR-455-3p TG group compared to WT (P=0.016) and KO mice (P=0.0003) (Fig. 2C). Further, time spent in the target quadrant that contains the platform was significantly increased in miR-455-3p TG mice relative to KO (P=0.003) and WT mice (P=0.015) (Fig. 2D). These observations indicate that the overexpressed miR-455-3p improve the spatial learning and memory in mice. The WT mice performed better than KO mice indicate that depletion of miR-455-3p impair the memory and cognitive function in mice.

**Figure 2.**
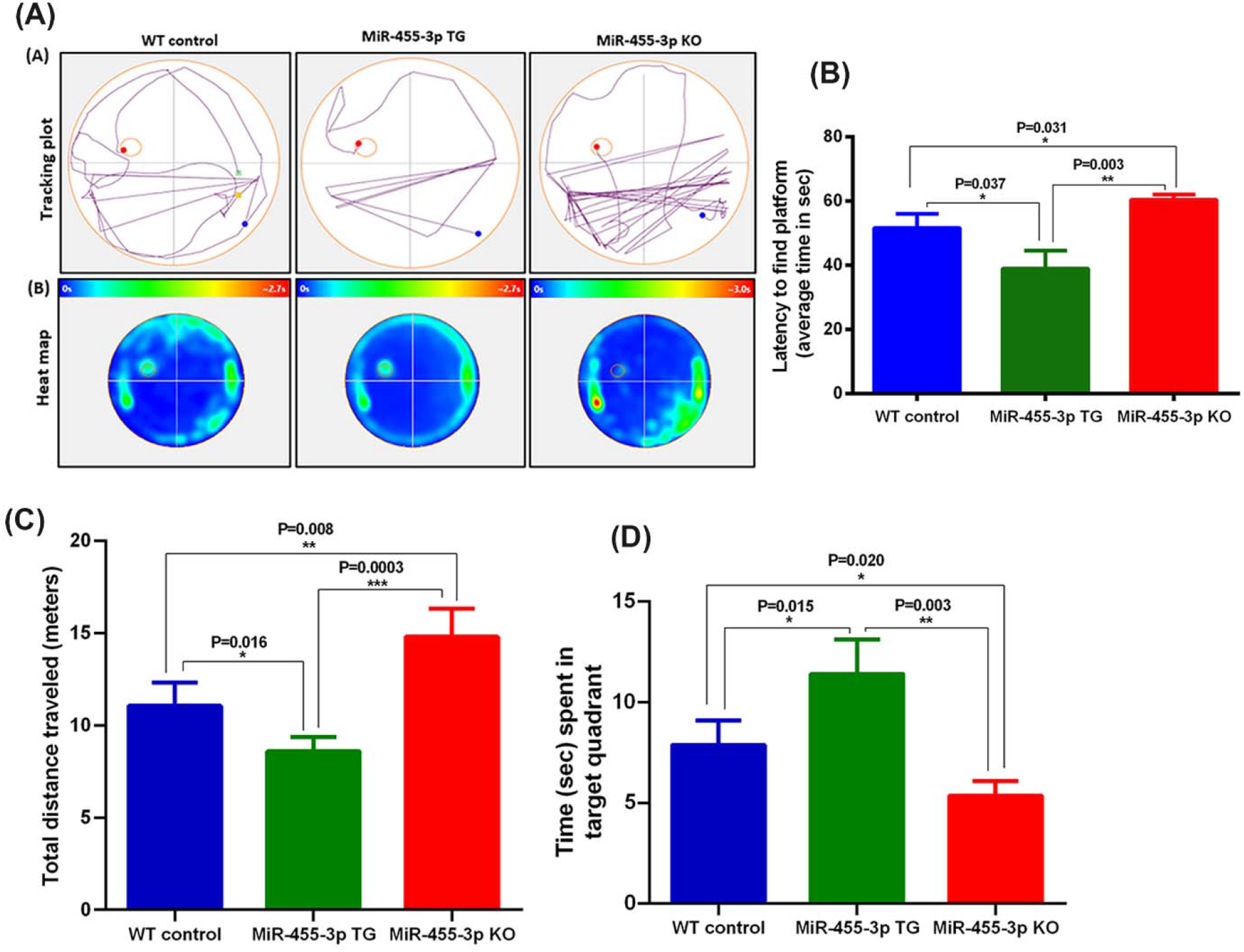
**A-** Mice cognitive behavior analysis using Morris Water Maze (MWM) test. (A) Representative mice body tracking plots showing mouse movement to find the platform in WT control, TG and KO mice. (B) Average heat map of MWM trials in WT (n=15), TG (n=15) and KO (n=15) mice showing mice locations during the test to find the platform. **B-** Latency to find the platform by WT mice, miR-455-3p TG and miR-455-3p KO mice. Average time (in seconds) to find the platform was significantly lower in TG mice relative to WT (P=0.037) and KO mice (P=0.003). KO mice spent more time to find the platform compared to WT mice. **C-** Total distance travelled (in meters) to find the platform by WT, TG and KO mice. The TG mice travelled significantly minimum distance compared to WT (P=0.016) and KO (P=0.003) while KO mice traveled maximum distance to find the platform compared to WT (P=0.008). **D-** Average time spent (in sec) in the target quadrant that having the platform by WT, TG and KO mice. TG mice spent significantly maximum time in target quadrant compared to WT (P=0.015) and KO mice (P=0.003). While KO mice spend minimum time in the target quadrant compared to WT mice (P=0.020).

### Impact of miR-455-3p on mRNA levels

To determine mRNA levels of mitochondrial biogenesis, dynamics and synaptic genes, we performed the qRT-PCR analysis of all mitochondrial biogenesis, dynamics and synaptic genes in WT, TG and KO mice. The TG mice showed the increased expression of biogenesis genes (PGC1α, Nrf1, Nrf2, TFAM), fusion proteins (Mfn1, Mfn2 and OPA1), synaptic genes (SNAP25, PSD95 and MAP2) and reduced expression of fission genes (Drp1 and Fis1) in TG mice relative to WT and KO mice (SI Fig. 5). While in KO mice, expressions of PGC1α, Nrf1, Nrf2, TFAM, Mfn1, Mfn2, OPA1, SNAP25, PSD95 and MAP2 were reduced compared to TG mice, and Drp1 and Fis1 expression levels were increased relative to TG mice. These results further validate the positive roles of miR-455-3p in regulation of mitochondrial biogenesis, dynamics and synaptic genes.

### Impact of miR-455-3p on protein levels-immunoblotting analysis

To determine the impact of miR-455-3p on mitochondrial biogenesis and dynamics activity, we analyzed the levels of mitochondrial biogenesis proteins (PGC1α, Nrf1, Nrf2 and TFAM) mitochondrial dynamics proteins (Drp1, Fis1, Mfn1, Mfn2 and Opa1) in WT, TG and KO mice brain tissues.

Fig. 3A, showed the representative immunoblots and Fig. 3B densitometry analysis. Our densitometry analysis showed the significantly increased levels of all four biogenesis proteins in TG mice relative to WT and KO mice (Fig. 3B). In KO mice, mitochondrial biogenesis protein levels were decreased significantly compared to WT control mice.

**Figure 3.**
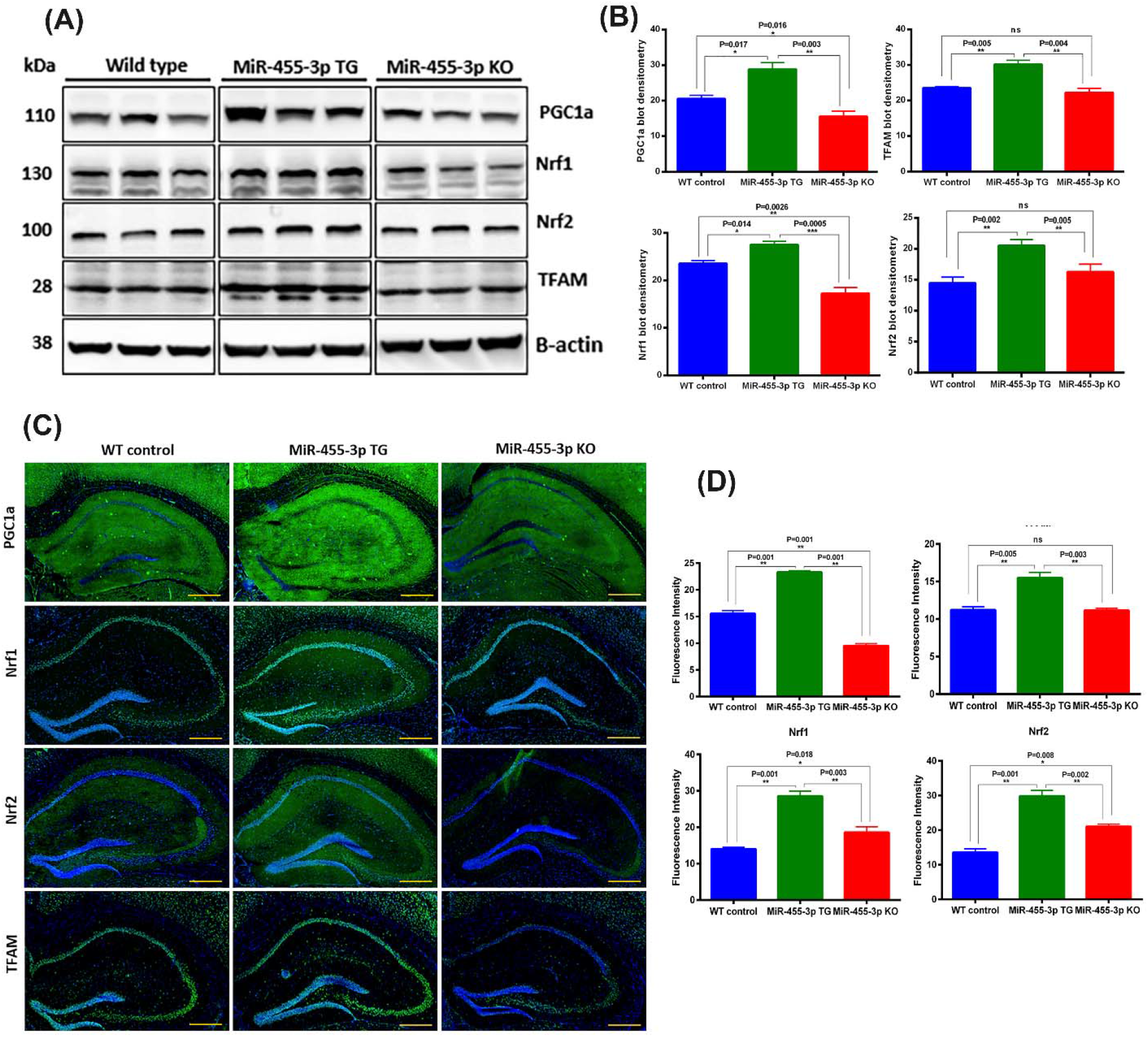
**A-** Immunoblot analysis of key mitochondrial biogenesis proteins in WT, TG and KO mice. Immunoblots for PGC1α, Nrf1, Nrf2, TFAM and B-actin proteins in representative animals WT (n=5), TG (n=5) and (n=5). **B-** Densitometry analysis for mitochondrial biogenesis proteins (PGC1α, Nrf1, Nrf2 and TFAM) in WT (n=5), TG (n=5) and KO (n=5) mice. All four biogenesis proteins increased significantly with different significant levels in TG mice relative to WT and KO mice. **C-** Immunostaining analysis of key mitochondrial biogenesis proteins in WT (n=3), TG (n=3) and KO (n=3) mice. Representative Immunostaining images for PGC1α, Nrf1, Nrf2 and TFAM proteins in WT, TG and KO mice at (10X magnification, 1mm scale bar). All four biogenesis proteins showed increased expression intensities in TG mice relative to WT and KO mice. **D-** Fluorescence intensity quantification for mitochondrial biogenesis proteins (PGC1α, Nrf1, Nrf2 and TFAM in WT, TG and KO mice. All four biogenesis proteins increased significantly with different significant levels in TG mice relative to WT and KO mice.

Next, we determined the protein levels of mitochondrial dynamics (fission proteins-Drp1 and Fis1) and (fusion proteins-Mfn1, Mfn2 and Opa1) in WT, TG and KO mice (SI Fig. 6). The densitometry analysis showed that mitochondrial fissions protein Fis1 was reduced significantly in TG mice relative to WT (P=0.043) and KO (P=0.012) mice, but Drp1 did not show any significant changes (SI Fig. 7). The mitochondrial fusion protein Mfn1 showed the significantly increased in TG mice relative to WT (P=0.004) and KO mice (P=0.0003). The Mfn2 protein levels did not showed significant changes in TG mice though it was significantly reduced (P=0.0063) in KO mice relative to WT mice. Further, Opa1 levels were also increase significantly in TG mice relative to KO mice (P=0.005).

To understand the impact of miR-455-3p on synaptic activity, we also assessed the levels of synaptic proteins- SNAP25, PSD95 and MAP2 in WT, TG and KO mice. The SI Fig. 8, shows representative immunoblots of SNAP25, PSD95 and MAP2 proteins in WT, TG and KO mice. Densitometry analysis revealed significantly increased levels of SNAP25, PSD95 and MAP2 proteins in TG mice compared to WT mice (SI Fig. 9).

### Impact of miR-455-3p on protein levels-immunostaining analysis

To understand the localization of mitochondrial biogenesis proteins, we performed the immunostaining analysis, focusing on hippocampal and cortical regions of WT, TG and KO mice (Fig. 3C). Fluorescence intensity quantification showed the significantly increased fluorescence intensities of all four proteins PGC1α, Nrf1, Nrf2 and TFAM in TG mice relative to WT and KO mice (Fig. 3D).

We also performed the immunostaining analysis of mitochondrial dynamics proteins in WT, TG and KO mice brains (SI Fig. 10). Most of the proteins showed the similar trend as seen by immunoblotting analysis. The fluorescence intensities of Drp1 fission protein were found to be significantly increased in KO mice relative to TG (P=0.021) and WT mice (P=0.002). The fluorescence intensity of all fusion proteins Mfn1, Mfn2 and OPA1 were found to be significantly increased in TG mice relative to TG and KO mice (SI Fig. 11). These results showed that miR-455-3p overexpression positively regulate the mitochondrial structural and functional activities.

The immunostaining data of synaptic proteins also showed increased levels of SNAP25, PSD95 and MAP2 in TG mice relative to WT and KO mice (SI Fig. 12, and 13). These results again confirmed that overexpression of miR-455-3p induces the synaptic proteins and could improve synaptic activities.

### Impact of miR-455-3p on brain cells populations

To determine the impact of miR-455-3p on brain cell specific population and differentiation- neuron, astrocytes and microglia, brain sections were immuno-stained using cell type-specific markers - NeuN, GFAP and Iba1 respectively. SI Fig. 14 showed the representative immunostaining images of neurons, astrocytes and microglia in WT, TG and KO mice. Fluorescence intensity quantification showed the significantly increased expression intensity of NeuN in TG mice relative to WT (P=0.002) and KO mice (P=0.0003) (SI Fig. 15). The KO mice showed significantly reduced levels of NeuN relative to WT mice (P=0.013). The astrocyte population (GFAP expression) was significantly reduced in TG mice relative to both WT (P=0.0004) and KO mice (P=0.005). Whereas astrocyte and microglia populations were found to be increased significantly in KO mice relative to TG (P=0.012) and WT mice (P=0.027) (SI Fig. 15). All together, these observations confirmed that miR-455-3p overexpression increased the neuronal populations and knockdown of miR-455-3p enhances astrocyte and microglial activity.

### Impact of miR-455-3p on dendritic spine density

To determine the impact of miR-455-3p on dendritic morphology i.e., length and spines number, we quantified dendritic length and number of spines using Golgi-Cox staining in the hippocampi of 12 months old WT, TG and KO mice. We compared the data within three groups of animals. The Fig. 4A showed the representative images of dendrites in WT, TG and KO mouse brains covering both cortex and hippocampus areas at 4X, 10X, 20X and 60X magnifications. Hippocampal neurons from TG mice showed the significant visual difference with dense and elongated dendrites compared to both WT and KO mice. The measurement of dendritic length showed significant difference in TG mice relative to WT (P=0.0005) and KO mice (P<0.0001) (Fig. 4B).

**Figure 4.**
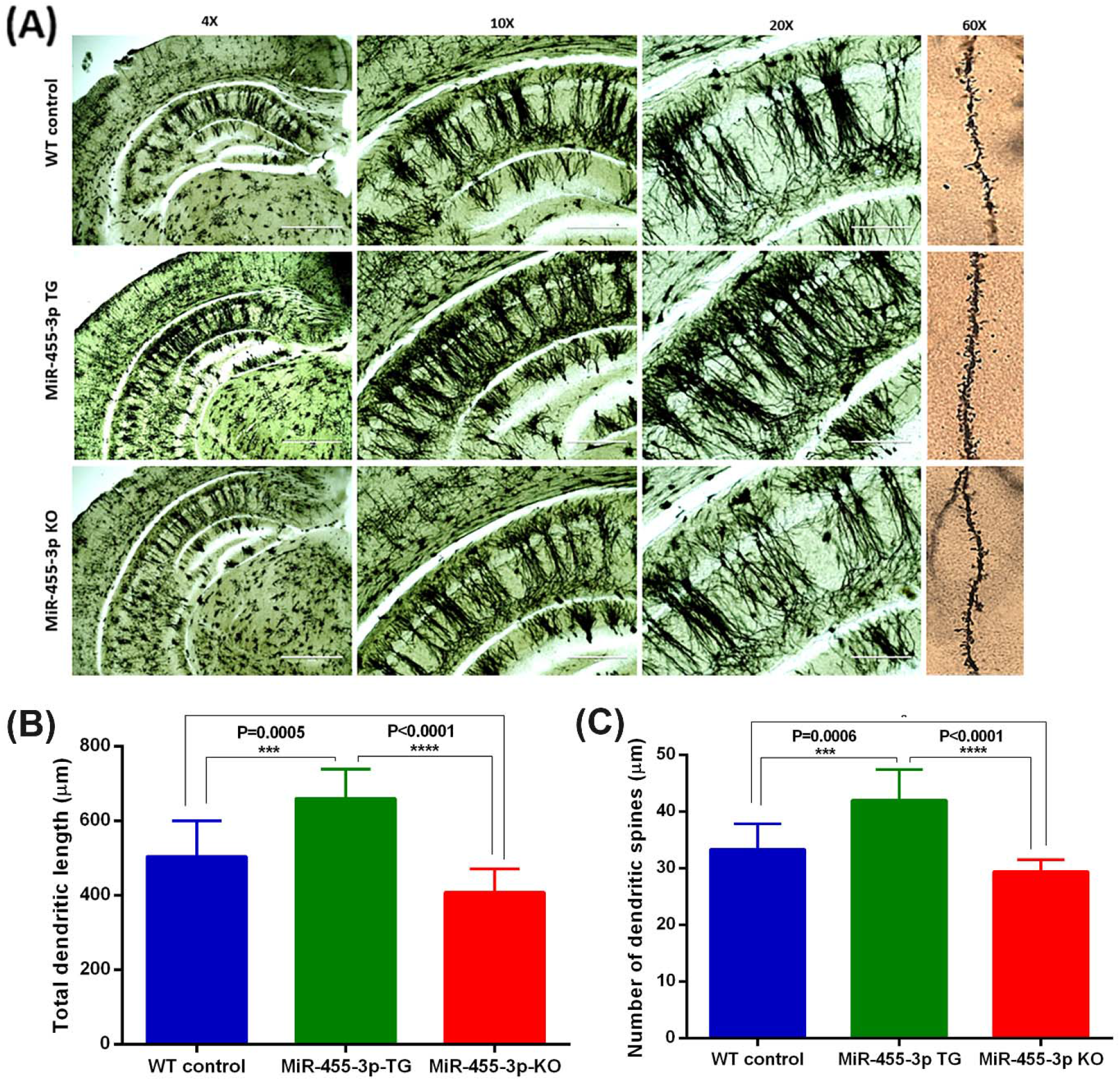
**A-** Dendritic spine density analysis of hippocampal neurons in WT, TG and KO mice. Representative Golgi-Cox staining images of dendrites in WT (n=3), TG (n=3) and KO (n=3) mice brains showing cortex and hippocampus areas at 4X, 10X, 20X and 60X magnifications (1 mm scale). **B-** Quantification of dendritic length (μm) in WT, TG and KO mice. The total dendritic length was significantly increased in TG mice relative to WT (P=0.0005) and KO mice (P<0.0001). In KO mice dendritic length was reduced significantly relative to normal WT mice (P=0.032). **C-** Quantification of dendritic spines numbers in WT, TG and KO mice. The average number of dendritic spines were significantly increased in TG mice relative to WT (P=0.0006) and KO mice (P<0.0001). In KO mice dendritic spine numbers were reduced significantly relative to normal WT mice (P=0.046).

Further, we quantified the number of dendritic spines using high magnification (60X) images in WT, TG and KO mice as showed in Fig. 4A. The number of dendritic spines were found to be significantly increased in TG mice compared to WT (P=0.0006) and KO mice (P<0.0001) (Fig. 4C). Interestingly, we also find the significant differences in the dendritic numbers and lengths in KO mice compared to WT mice. These observations indicate that miR-455-3p overexpression enhances both dendritic length and number of dendritic spines and reduced in knockout miR-455-3p mice. These results confirmed the significant positive impact of miR-455-3p on hippocampal neurons, dendritic morphology and quality.

### Impact of miR-455-3p on mitochondrial morphology (number and length)

To determine the impact of miR-455-3p on mitochondrial number and length, we used transmission electron microscopy on cortical and hippocampal tissues from 12-month-old WT, TG and KO mice. We analyzed the data in terms of mitochondrial number and length in all three groups of mice. Fig. 5A showed representative images of mitochondrial morphology in cortical and hippocampal areas of WT, TG and KO mice brains. Fig. 5B showed significantly reduced mitochondrial number in cortex of TG mice relative to WT (P=0.013) and KO mice (P=0.0001) (Fig. 5B). Interestingly, KO mice also showed the significantly increased mitochondrial (fragmentation) numbers (P=0.002) relative to WT mice.

**Figure 5.**
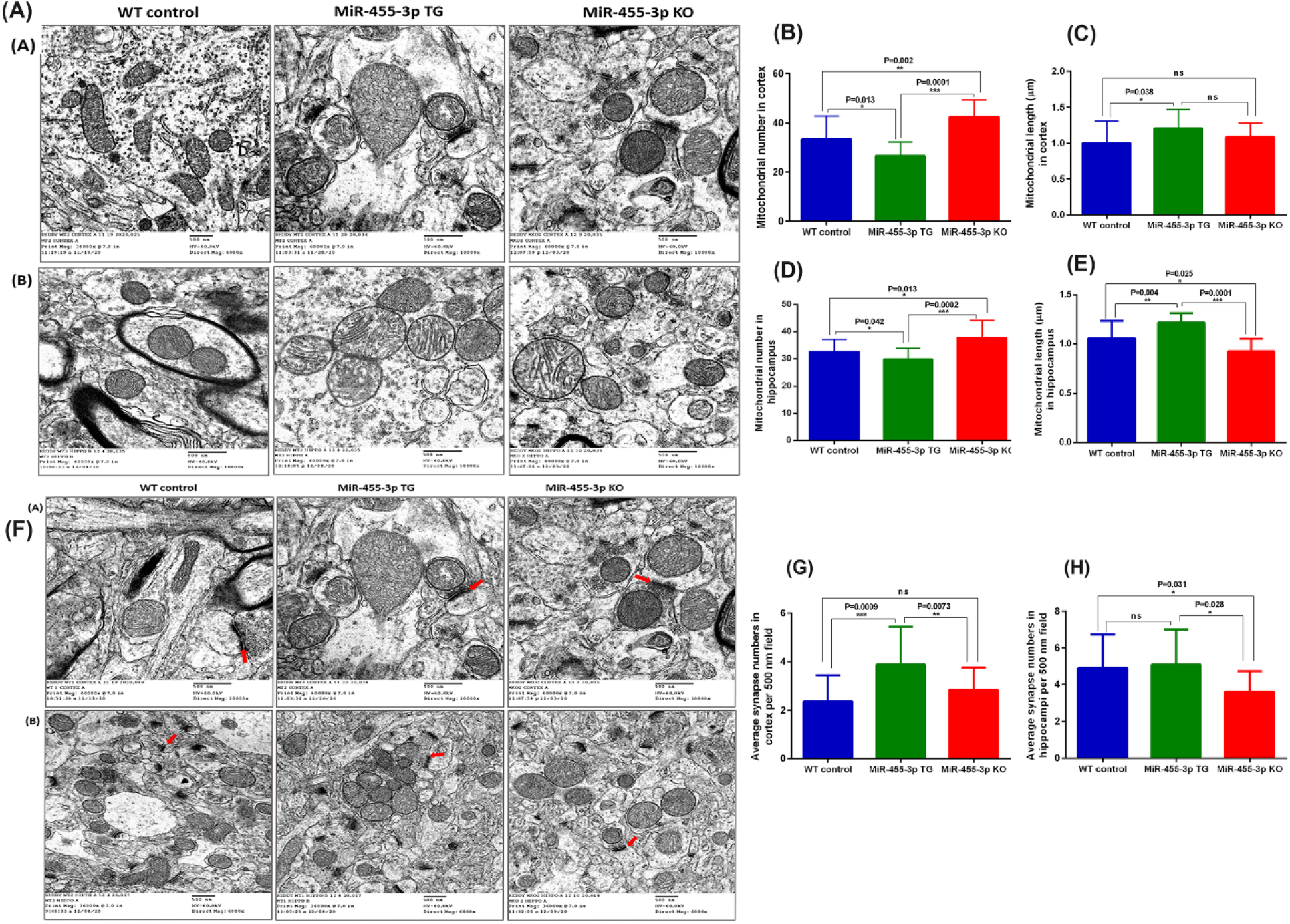
**A-** Transmission electron microscopic analysis of mitochondrial morphology. Representative TEM images of (A) Cortex and (B) Hippocampus in WT (n=3), TG (n=3) and KO (n=3) mice brains showing mitochondrial organization at 500 nm magnifications. **B-** Quantification of mitochondrial numbers in WT, TG and KO mice cortex. The mitochondrial numbers were significantly lower in TG mice relative to WT (P=0.013) and KO mice (P=0.0001). Even WT mice has significantly lower numbers than KO mice (P=0.002). **C-** Quantification of mitochondrial length (μm) in WT, TG and KO mice cortex. The mitochondrial length was significantly increased in TG mice relative to WT (P=0.038) mice. The KO mice did not show significant variation in mitochondrial length relative to WT and TG mice. **D-** Quantification of mitochondrial numbers in WT, TG and KO mice hippocampus. The mitochondrial numbers were significantly lower in TG mice relative to WT (P=0.042) and KO mice (P=0.0002). Even WT mice has significantly lower numbers than KO mice (P=0.013). **E-** Quantification of mitochondrial length (μm) in WT, TG and KO mice hippocampus. The mitochondrial length was significantly increased in TG mice relative to WT (P=0.004) and KO mice (P=0.0001). Even WT mice showed significantly increased mitochondrial length compared to KO mice (P=0.025). **F-** Representative TEM images of cortex and hippocampus in WT (n=3), TG (n=3) and KO (n=3) mice brains showing synapse assembly and synapse number at 500 nm magnifications. Red arrow showing the synaptic cleft. **G-** Quantification of synapse numbers in WT, TG and KO mice cortex. The synapse numbers were found to be significantly increased in TG mice relative to WT (P=0.0009) and KO mice (P=0073). WT and KO mice did not showed significant changes in synapse numbers. **H-** Quantification of synapse numbers in WT, TG and KO mice hippocampi. The synapse numbers were found to be significantly increased in TG mice relative to KO mice (P=028). In KO mice synapse numbers decreased significantly relative to WT mice (P=0.031). The WT and TG mice did not showed significant changes in synapse numbers.

On the other hand, mitochondrial length was significantly increased in TG mice relative to WT mice (P=0.038), however we did not see the any significant difference in mitochondrial length in KO mice compared to TG mice (Fig. 5C).

Next, we examined the mitochondrial morphology in WT, TG and KO mice hippocampus area (Fig. 5A). We did see the significant difference in the mitochondrial numbers among WT vs TG vs KO mice hippocampal area. Fig. 5D, shows significantly reduced mitochondrial number in TG mice relative to WT (P=0.042) and KO mice (P=0.0002). Whereas mitochondrial length was significantly increased in TG mice relative to WT (P=0.004) and KO mice (P=0.0001) (Fig. 5E).

The KO mice showed the poor mitochondrial quality in terms of mitochondrial fragmentation compared to WT and TG mice. These observations confirm that high levels of miR-455-3p improve the mitochondrial morphology, in terms of number and length in both cortex and hippocampi.

### Impact of miR-455-3p on synapse numbers

We also examined the impact of miR-455-3p on synapse organization and numbers in both cortex and hippocampus of WT, TG and KO mice. The Fig. 5F showed the synapse location and organization in WT, TG and KO mice cortex, the red arrow showed the synaptic cleft. As shown in Fig. 5G, the average synapse numbers were significantly increased in TG mice cortex relative to both WT (P=0.0009) and KO mice (P=0.0073). The Fig. 5F, showed the synapse locations and numbers in WT, TG and KO mice hippocampus. Quantification showed the significant increased synapse numbers in TG mice relative to KO (P=0.028) but we did not find the significant difference in TG vs WT mice (Fig 5H). These observations confirm that miR-455-3p might also involve in synapse assembly and junction formation.

## Discussion

Mouse models are the most important research tools to study the molecular basis of diseases and to test potential therapeutic interventions. Several AD mouse models (TG, KO, Knock-in) are used to understand disease process from birth to terminal stages and also therapeutic drugs. And also, AD mouse models used to test potential anti-AD agents and to study the mutational impact of protective human genes/proteins on AD mice via genetic crossing with Tg and KO mouse models [29,30].

MiRNA mouse models are emerging tools to extensively study the protective and/or deleterious effects of miRNAs in human diseases [31,32]. Since, single miRNA, for example miR-455-3p is implicated in multiple human diseases such as cancers and chondrogenesis via modulating a bunch of potential target proteins [21]. Recently, our lab discovered the properties of miR-455-3p in AD [21]. Based on the critical roles of miR-455-3p in AD and anti-AD characteristics, it was warranted to develop miR-455-3p mouse models. Other than AD, generation of miR-455-3p mouse models could be useful for miR-455-3p dependent diseases also.

MiRNAs molecules has been identified as strong modulators of human genes and/or proteins endogenously at genetic and epigenetic levels [33]. Several miRNAs have been studied previously in AD that showed promising results against AD genes/proteins via modulation of key AD related genes [13–20]. However, most of miRNA(s) studies did not go longer in depth analysis due to lack of suitable mouse model and intervention mode. Quite a few studies showed the miRNAs intervention using either direct oral/nasal miRNAs complex administration or by injections [34]. Again, these studies had several limitations, 1) dose and route of miRNA administration, 2) miRNA complex stability, 3) immune reactivity of miRNAs encapsulated particles, 4) half-life of exogenous miRNA complex inside the body and 5) degree of effectiveness. To overcome these challenges and technical barriers the generation of miRNA mouse model is ideal and provides detailed molecular mechanisms and therapeutic potentials of a particular miRNA. Based on the previous miR-455-3p studies [23–26] and its promising anti-AD properties [21], the current study was attempted to develop miR-455-3p TG and KO mouse models.

In the current study, miR-455-3p TG and KO models were successfully generated by using pronuclear injection of miR-455-3p transgene to mouse embryo and CRISPER/Cas9 knockout techniques. Both TG and KO mice were characterized by genotyping, sequencing, miR-455-3p qRT-PCR and immunoblotting studies. Both mouse models displayed normal phenotypic characteristics and we did not observe any serious deformities due to miR-455-3p transgenes insertion and depletion.

### Lifespan extension

We followed TG, KO and non-transgenic and non-KO WT mice from birth to terminal stages of life, and our survival data revealed that TG mice lived 5 months longer than age-matched non-transgenic WT mice; on the other hand, KO mice survived 4 months shorter than their non-transgenic WT counter parts. These observations indicate overexpression of miR-455-3p extend overall lifespan, in other words, miR-455-3p has protective properties of brain and peripheral organs. As described, our current study focused on brain, it is worthwhile to study other organs, such, heart, lung, liver, kidney, and skeletal muscle, in order to understand protective properties, how miR-455-3p protects different organs. Based on our current study, endogenous full-length APP levels were reduced in TG-miR-455-3p mice, and these were increased in KO mice, it is possible that reduction of full-length is protective to brain and other peripheral organs.

If miR-455-3p is capable of reducing full-length APP and its toxicities in mice, it is important to cross miR-455-3p TG and KO with mouse models of AD, particularly mice with intact 3’ UTR, say newly developed humanized Ab-KI mice. The resulting double mutant mice TG miR-455-3p X hAb-KI predicted to live longer & show reduced cognitive decline and APP and its c-terminal derivative toxicities of mitochondria and synaptic activities. On the other hand, double mutant KO miR-455-3p X hAb-KI mice expected to accelerate AD pathologies, increased cognitive decline and reduced lifespan.

### Cognitive behavior

The main physiological hallmark of AD pathogenesis is cognitive deficits, so we examined the miR-455-3p effect on mice cognitive behavioral activities. The TG mice showed better cognitive learning and memory compared to age matched WT mice in MWM test. We also observed that KO mice did not perform well on MWM test. These results showed that high endogenous miR-455-3p improves cognitive functions. To understand the molecular mechanism for cognition improvement by miR-455-3p, we studied the synaptic proteins, synapse assembly organization, synapse numbers and dendritic spine density of cortical and hippocampal neurons. Since healthy and active synapse are the key for a typical synaptic activity [35–38].

### Synaptic and dendritic activity

The TG mice demonstrated increased levels of SNAP25, PSD95 and MAP2 proteins and essentially increased synapse numbers in both cortex and hippocampus. In addition, the dendritic spine density of hippocampal neurons was also increased in miR-455-3p TG mice. However, it is not clear how miR-455-3p upregulation enhances synaptic proteins and synaptic activity. It is well established that cognitive function was improved by increased synaptic proteins [37] and by upregulation of PGC1α [39]. Recent studies showed the improvement of cognitive function mediated by miR-455-3p upregulation via suppression of Histone deacetylase 2 (HDAC) and EphA4 [40,41]. High levels of miR-455-3p reduces the neurotoxicity by suppressing the EphA4 expression in hippocampal neurons [41]. Other study found the relevance of miR-455-3p in restoring neurological function and improving memory deficits in traumatic brain injury (TBI) [40]. Actually, cinnamic acid as a treatment of TBI increases the expression of miR-455-3p which intern suppressed the HDAC2 and reduce memory impairments. Our findings are in the line with these published reports and support our results

### Cell type analysis

We studied the impact of miR-455-3p on brain cells differentiation. Interestingly, neuronal populations were found to be increased in TG mice as shown by NeuN staining. Even increased levels of other neuronal marker MAP2, in TG mice again confirmed the increased neuronal population by miR-455-3p over expression. Interestingly, astrocyte population was significantly reduced in TG mice compared to both WT and KO. On the other side, knockdown of miR-455-3p showed reduced levels of neuronal staining and KO mice showed the increased microglia and astrocyte expression.

### Mitochondrial structure and function

We also examined the miR-455-3p impact on mitochondrial structure and function. Since mitochondrial structural and functional abnormalities are primarily involved in disease progression and synaptic dysfunction in AD [42,43]. As a parameter of mitochondrial quality, the TEM analysis for the evaluation of mitochondrial number and length showed improved mitochondrial organization in miR-455-3p TG mice. The improved mitochondrial quality was correlated with the levels of mitochondrial biogenesis proteins in TG mice. All four biogenesis proteins (PGC1α, NRF1, NRF2, and TFAM) were elevated in TG mice and the levels of dynamics proteins (DRP1 and FIS1, OPA1, Mfn1, and Mfn2) were positively regulated by miR-455-3p. However, it is argued that how miR-455-3p regulate the mitochondrial functions. A well-known mechanism is modulation of HIF1a by miR-455-3p [22]. On the other side HIf1a interaction with PGC1α is well established in mitochondrial function [44,45]. Therefore, miR-455-3p mediated HIF1a and PGC1α are the key drivers that controls mitochondrial function in TG mice.

In summary, based on the miR-455-3p properties, current study is focused on the generation and characterization of miR-455-3p TG and KO mouse models to understand the molecular mechanism of miR-455-3p in mice survival, APP protein, brain cells proliferation, synapse assembly formation, mitochondrial function, synaptic activity and mice cognitive behaviors. MiR-455-3p TG mice showed the improved life span, neuronal growth, synapse, cognitive function and overall mitochondrial and synaptic functions. While miR-455-3p KO mice exhibited reduced life span, neuronal growth, memory, mitochondrial and synaptic function. In conclusion, miR-455-3p exhibits positive impacts on neuronal activity in several ways and improve overall brain functions. Further, genetic crossing study is needed to understand the protective effect of miR-455-3p on the mutant APP proteins using hAb-KI and APP Tg mice lines. Additionally, these mouse models could be ideal tool understand the molecular mechanism of miR-455-3p in AD and other diseases.

## Acknowledgements

The authors would like to thank NIH for funding various projects - R01AG042178, R01AG47812, R01NS105473, AG060767, AG069333, and AG066347 to (P.H.R) and K99AG065645 to (S.K.).

