## Supplemental Table 2 for "Novel microRNA-455-3p mouse models to study Alzheimer’s disease pathogenesis"

**SI Table 2. Summary of antibody dilutions and conditions used in the immunoblotting analysis of APP, mitochondrial biogenesis, mitochondrial dynamics and synaptic proteins in WT, miR-455-3p TG and miR-455-3p KO mice**

| **Marker** | **Primary Antibody – Species and Dilution** | **Purchased from Company, City & State** | **Secondary Antibody, Dilution** | **Purchased from Company, City & State** |
| --- | --- | --- | --- | --- |
| PSD95 | Rabbit Monoclonal 1:300 | Novus Biological,  Littleton, CO | Donkey Anti-rabbit HRP 1:10,000 | GE Healthcare Amersham,  Piscataway, NJ |
| SNAP25 | Rabbit  Polyclonal  1:500 | Novus Biological,  Littleton, CO | Donkey Anti-rabbit  HRO 1:10,000 | GE Healthcare Amersham,  Piscataway, NJ |
| MAP2 | Mouse  Monoclonal  1:500 | Invitrogen  Waltham, MA | Sheep Anti-mouse HRP 1:10,000 | GE Healthcare Amersham,  Piscataway, NJ |
| PGC1α | Rabbit Polyclonal 1:500 | Novus Biological,  Littleton, CO | Donkey Anti-rabbit HRP 1:10,000 | GE Healthcare Amersham,  Piscataway, NJ |
| Nrf1 | Rabbit Polyclonal 1:300 | Novus Biological,  Littleton, CO | Donkey Anti-rabbit HRP 1:10,000 | GE Healthcare Amersham,  Piscataway, NJ |
| Nrf2 | Rabbit Polyclonal 1:300 | Novus Biological,  Littleton, CO | Donkey Anti-rabbit HRP 1:10,000 | GE Healthcare Amersham,  Piscataway, NJ |
| TFAM | Rabbit Polyclonal 1:300 | Novus Biological,  Littleton, CO | Donkey Anti-rabbit HRP 1:10,000 | GE Healthcare Amersham,  Piscataway, NJ |
| Mfn1 | Rabbit Polyclonal 1:300 | Novus Biological,  Littleton, CO | Donkey Anti-rabbit HRP 1:10,000 | GE Healthcare Amersham,  Piscataway, NJ |
| Mfn2 | Rabbit Polyclonal 1:300 | Novus Biological,  Littleton, CO | Donkey Anti-rabbit HRP 1:10,000 | GE Healthcare Amersham,  Piscataway, NJ |
| OPA1 | Rabbit Polyclonal 1:300 | Novus Biological,  Littleton, CO | Donkey Anti-rabbit HRP 1:10,000 | GE Healthcare Amersham,  Piscataway, NJ |
| DRP1 | Rabbit Polyclonal 1:300 | Novus Biological,  Littleton, CO | Donkey Anti-rabbit HRP 1:10,000 | GE Healthcare Amersham,  Piscataway, NJ |
| FIS1 | Rabbit Polyclonal 1:300 | Novus Biological,  Littleton, CO | Donkey Anti-rabbit HRP 1:10,000 | GE Healthcare Amersham,  Piscataway, NJ |
| B-actin | Mouse Monoclonal 1:500 | Sigma-Aldrich,  St Luis, MO | Sheep Anti-mouse HRP 1:10,000 | GE Healthcare Amersham,  Piscataway, NJ |
| Mouse endogenous APP | Mouse Monoclonal 1:500 | Biolegend,  San Diego, CA | Sheep Anti-mouse HRP 1:10,000 | GE Healthcare Amersham,  Piscataway, NJ |
| NeuN | Rabbit Polyclonal 1:200 | Abcam, USA | Donkey Anti-rabbit HRP 1:1,000 | GE Healthcare Amersham,  Piscataway, NJ |
| GFAP | Mouse Monoclonal 1:300 | Novus Biological,  Littleton, CO | Donkey Anti-rabbit HRP 1:1,000 | GE Healthcare Amersham,  Piscataway, NJ |
| Iba1 | Rabbit Polyclonal 1:200 | Cell signaling,  USA | Donkey Anti-rabbit HRP 1:1,000 | GE Healthcare Amersham,  Piscataway, NJ |
