## Supplemental Table 1 for "Novel microRNA-455-3p mouse models to study Alzheimer’s disease pathogenesis"

**SI Table 1. Summary of qRT-PCR oligonucleotide primers used in measuring miRNA and mRNA expression in WT, miR-455-3p TG and miR-455-3p KO mice**

| **miRNA** | **DNA Sequence (5’ to 3’)** | **Base pairs** |
| --- | --- | --- |
| miR-455-3p | Forward primer GCAGTCCATGGGCATATACAC | 68 |
| snoRNA-202 | Forward primer AGTACTTTTGAACCCTTTTCCA | 69 |
| APP | Forward primer TGGAGGTACCCACTGATGGT  Reverse primer TGTGCATGTTCAGTCTGCCA | 81 |
| PGC1α | Forward primer GCAGTCGCAACATGCTCAAG  Reverse primer GGGAACCCTTGGGGTCATTT | 83 |
| NRF1 | Forward primer AGAAACGGAAACGGCCTCAT  Reverse primer CATCCAACGTGGCTCTGAGT | 96 |
| NRF2 | Forward primer ATGGAGCAAGTTTGGCAGGA  Reverse primer GCTGGGAACAGCGGTAGTAT | 96 |
| TFAM | Forward primer TCCACAGAACAGCTACCCAA  Reverse primer CCACAGGGCTGCAATTTTCC | 84 |
| PSD95 | Forward primer CTTCATCCTTGCTGGGGGTC  Reverse primer TTGCGGAGGTCAACACCATT | 90 |
| SNAP25 | Forward primer GTGGTCATCTGGTGGCTCTA  Reverse primer TTCAGCAAACGCCACTGAGA | 91 |
| MAP2 | Forward Primer AGGAAGCCCAACACAAGGAC  Reverse Primer TTTGTTCTGAGGCTGGCGAT | 102 |
| DRP1 | Forward Primer ATGCCAGCAAGTCCACAGAA  Reverse Primer TGTTCTCGGGCAGACAGTTT | 86 |
| FIS1 | Forward Primer CAAAGAGGAACAGCGGGACT  Reverse Primer ACAGCCCTCGCACATACTTT | 95 |
| OPA1 | Forward Primer ACCTTGCCAGTTTAGCTCCC  Reverse Primer TTGGGACCTGCAGTGAAGAA | 82 |
| Mfn1 | Forward Primer GCAGACAGCACATGGAGAGA Reverse Primer GATCCGATTCCGAGCTTCCG | 83 |
| Mfn2 | Forward Primer TGCACCGCCATATAGAGGAAG  Reverse Primer TCTGCAGTGAACTGGCAATG | 78 |
