## Supplemental information for "Novel microRNA-455-3p mouse models to study Alzheimer’s disease pathogenesis"

### 1. Transgenic Products

#### Product 1:

| **Name of Injected DNA** | pRP[Exp]-U6>hsa-miR-455-3p-CAG-EGFP | | |
| --- | --- | --- | --- |
| **Mouse Strain** | C57BL/6×C57BL/6 | | |
| **Date of Birth** | 06-22-2018 | | |
| **Description** | Total: 9 pups | | |
|  | pRP[Exp]-U6>hsa-miR-455-3p-CAG-EGFP: 56-64 (From founder #40) | | |
| **PCR-Positive Pups** | ♂ | - | pRP[Exp]-U6>hsa-miR-455-3p-CAG-EGFP: - |
|  | ♀ | 1 | pRP[Exp]-U6>hsa-miR-455-3p-CAG-EGFP: 63 |

**Product 2:**

| **Name of Injected DNA** | pRP[Exp]-U6>hsa-miR-455-3p-CAG-EGFP | | |
| --- | --- | --- | --- |
| **Mouse Strain** | C57BL/6×C57BL/6 | | |
| **Date of Birth** | 07-15-2018 | | |
| **Description** | Total: 9 pups | | |
|  | pRP[Exp]-U6>hsa-miR-455-3p-CAG-EGFP: 15-23 (From founder #40) | | |
| **PCR-Positive Pups** | ♂ | - | pRP[Exp]-U6>hsa-miR-455-3p-CAG-EGFP: - |
|  | ♀ | 1 | pRP[Exp]-U6>hsa-miR-455-3p-CAG-EGFP: 15 |

2. **PCR Conditions**

The pups were screened by the following PCR assay. Out of 18 pups screened, 2 were identified positive, which were then confirmed by the same PCR with recut samples.

#### Primers Used:

Transgene PCR primer F1: GTTCGGCTTCTGGCGTGTG Transgene PCR primer R1: TTCAGGGTCAGCTTGCCGTAG Annealing Temp: 60 ℃

#### Expected PCR Product:

Transgene PCR product size: 297 bp

#### Primers Used:

Transgene PCR primer F2: TCAAGATCCGCCACAACATCG Transgene PCR primer R2: CACTGCATTCTAGTTGTGGTTTGTC Annealing Temp: 60 ℃

#### Expected PCR Product:

Transgene PCR product size: 318 bp

Quote: TGMB-170927-AHL-01+TGMB-170927-AHL-01_SUP1

2 / 6

*2255 Martin Ave., Suite E. Santa Clara, CA 95050*

### 3. PCR Result

#### Note:

1) PCR was carried out in 25 µL volume for 35 cycles under standard conditions, with all two primers listed above added to each reaction.
2) Taq DNA polymerase used was P112-01.
3) Three controls used in PCR genotyping are:

- Water control: No DNA template added.
- Positive control: 400 ng of mouse genomic DNA spiked with an amount of transgene injection DNA that is equivalent to 5 copies of transgene per diploid mouse genome.
- Wildtype control: 400 ng of mouse genomic DNA.

#### Client should report any problems with the delivered mice and other materials to Cyagen within one week of receiving the shipment.

4. **PCR Conditions Attachment**
   4.1 **DNA Extraction**

- Method One:

We recommend that using TaKaRa MiniBEST Universal Genomic DNA Extraction kit (Ver.5.0_Code No. 9765) to gain high purity of genomic DNA.

Quote: TGMB-170927-AHL-01+TGMB-170927-AHL-01_SUP1

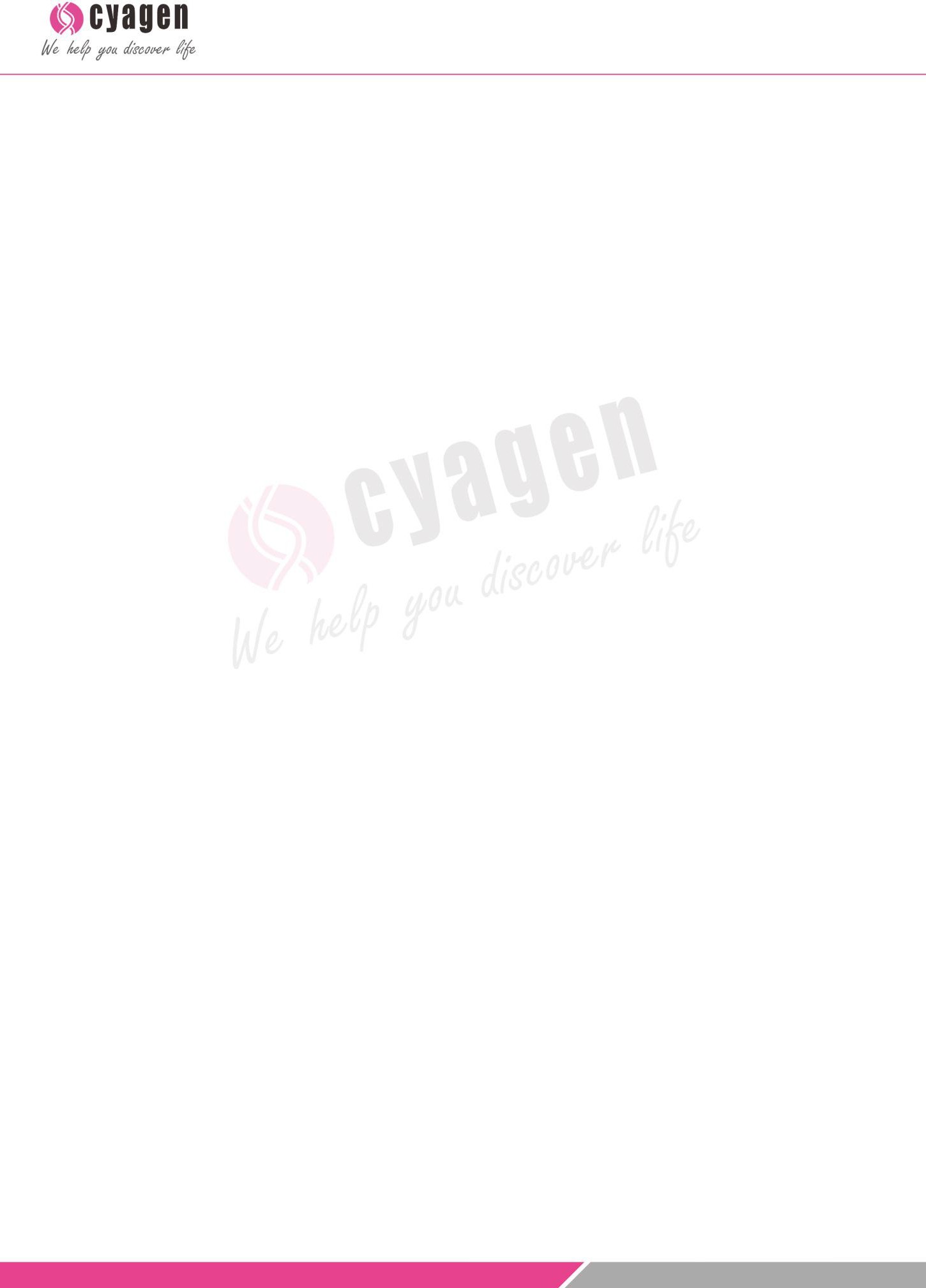

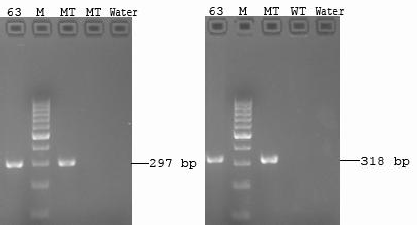

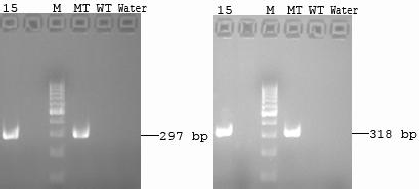

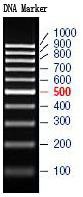

3 / 6

*2255 Martin Ave., Suite E. Santa Clara, CA 95050*

a. Add 180 μL of Buffer GL, 20 μL of Proteinase K and 10 μL of RNase A per tail piece (2-5 mm) in a microcentrifuge tube. Be careful not to cut too much tail.
b. Incubate the tube at 56 ℃ overnight.
c. Spin in microcentrifuge at 12,000 rpm for 2 minutes to remove impurities.
d. Add 200 μL Buffer GB and 200 μL absolute ethyl alcohol with sufficient mixing.
e. Place the spin Column in a collection tube. Apply the sample to the spin and centrifuge at 12,000 rpm for 2 min. Discard flow-through.
f. Add 500 μL Buffer WA to the spin column and centrifuge at 12,000 rpm for 1 min. Discard flow-through.
g. Add 700 μL Buffer WB to the spin column and centrifuge at 12,000 rpm for 1 min. Discard flow-through. (Note: Make sure the Buffer WB has been premixed with 100% ethanol. When adding Buffer WB, add to the tube wall to wash off the residual salt.)
h. Repeat step g.
i. Place the spin Column in a collection tube and centrifuge at 12,000 rpm for 2 min.
j. Place the spin Column in a new 1.5ml tube. Add 50~200 μL sterilized water or elution buffer to the center of the column membrane and let the column stand 5min. (Note: Heating sterilized water or elution buffer up to 65℃ can increase the yield of elution.)
k. To elute DNA, centrifuge the column at 12,000 rpm for 2 min. To increase the yield of DNA, add the flow-through and/or 50~200 μL sterilized water or elution buffer to the center of the spin column membrane and let the column stand 5 min. Centrifuge at 12,000 rpm for 2 min.
l. Quantify to genomic DNA. Eluted genomic DNA can be quantified by electrophoresis or electrophoresis.

- Method Two:

A low-cost and sample method to gain rough genomic DNA.

a. Add 100 μL of tail digestion buffer per tail piece (2-5 mm) in a microcentrifuge tube. Be careful not to cut too much tail.
b. Incubate the tube at 56 ℃ overnight.
c. Incubate the tube at 98 ℃ for 13 minutes to denature the Proteinase K.
d. Spin in microcentrifuge at top speed for 15 minutes. Use an aliquot of supernatant straight from the tube (1.5 μL in a 25 μL reaction) for PCR.

Final concentration of tail digestion buffer:

- 50 mM KCl
- 10 mM Tris-HCl (pH 9.0)
- 0.1 % Triton X-100
- 0.4 mg/mL Proteinase K

#### 4.2 PCR Mixture (primer concentration: 10μM ):

Component	x1

ddH_2_O	9.0 μl

Quote: TGMB-170927-AHL-01+TGMB-170927-AHL-01_SUP1

4 / 6

*2255 Martin Ave., Suite E. Santa Clara, CA 95050*

Product primer F	1.0 μl

Product primer R	1.0 μl

Premix Taq	12.5 μl

DNA	1.5 μl

Total	25	μl

#### 4.3 PCR Reaction Conditions:

Step	Temp.	Time	Cycles

Initial denaturation	94 °C	3 min

Denaturation	94 °C	30 s

Annealing	60 °C	35 s	35 x

Extension	72 °C	35 s

Additional extension	72 °C	5 min

#### 4.4 Relevant Reagents:

| **Trizma Hydrochloride Solution** | Sigma, Cat. No. T2663 |
| --- | --- |
| **Proteinase K** | Merck, Cat. No. MK539480 |
| **Triton X-100** | Sigma, T8787-50 mL |
| **2 × Taq Master Mix (Dye Plus)** | Vazyme, P112-01 |
| **Agarose** | BIOWEST AGAROSE, REGULAR |
| **DNA Marker** | Thermo Scientific GeneRuler 100 bp DNA Ladder #SM0242 |
| **0.5×TBE** | Tris Bio Basic Inc, TBO194-500g |
|  | EDTA Shanghai Sangon, 0105-500g |
|  | Boric Acid, Shanghai Sangon, 0588-500g |

Quote: TGMB-170927-AHL-01+TGMB-170927-AHL-01_SUP1

5 / 6

*x: 408-969-0336*
