## Supplemental information for "Novel microRNA-455-3p mouse models to study Alzheimer’s disease pathogenesis"

**Mouse Mir455 Knockout Project (CRISPR/Cas9)**

1. OBJECTIVE

To create a Mir455 3P knockout mouse model (C57BL/6) by CRISPR/Cas-mediated genome engineering.

2. PROJECT SUMMARY

- The mouse Mir455 gene (miRBase: MI0004679; Ensembl: ENSMUSG00000070102) is located on mouse chromosome 4.
- 3P of mouse Mir455 will be selected as target site (sequences shown on the next page).
- Two pairs of gRNA targeting vectors will be constructed and confirmed by sequencing.
- Cas9 mRNA, gRNA generated by *in vitro* transcription will be co-injected into fertilized eggs for KO mouse production.
- The pups will be genotyped by PCR followed by sequence analysis.

3. TARGETING STRATEGY

**3a. Schematic depiction of targeting strategy**

Genomic region of mouse Mir455 locus is diagrammed below (gene is oriented from left to right; total size is 82 bp).

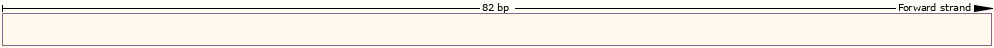

gRNA4

gRNA3

gRNA2

gRNA1

3’

5’

3P of mouse Mir455

5P of mouse Mir455

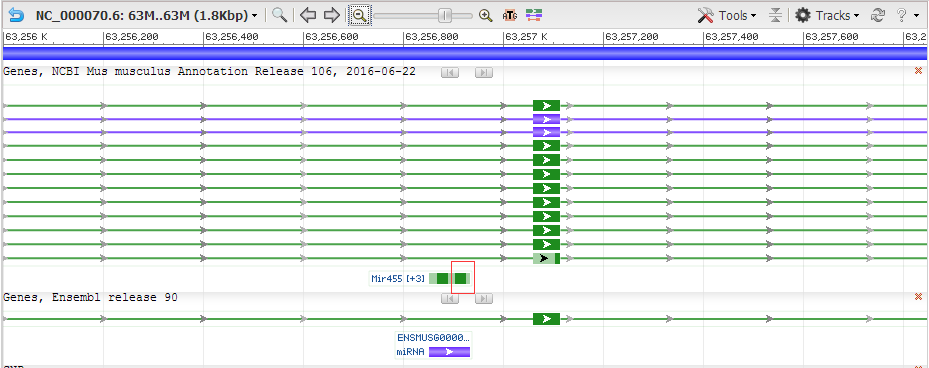

**3b. gRNA target sequence**

Pair 1

gRNA1 (matches reverse strand of gene): GTATATGCCCGTGGACTGCATGG

gRNA2 (matches forward strand of gene): CCTCAAGGTCTATGTCATCGAGG

Pair 2

gRNA3 (matches forward strand of gene): CGCAGCACCATGCAGTCCACGGG

gRNA4 (matches forward strand of gene): TCTATGTCATCGAGGACCCCTGG

**3c.The links of gRNA on VectorBuilder**

gRNA1: <https://www.vectorbuilder.com/vector/VB170929-1146nyr.html>

gRNA2: <https://www.vectorbuilder.com/vector/VB170929-1147pdz.html>

gRNA3: <https://www.vectorbuilder.com/vector/VB170929-1148dum.html>

gRNA4: <https://www.vectorbuilder.com/vector/VB170929-1150ugt.html>

**3d. Assay of CRISPR-induced mutation**

The target region of mouse Mir455 locus will be amplified by PCR with specific primers. PCR product will be sequenced to confirm targeting.

Primer sequence:

Mouse Mir455-pair1/pair2-F: CAGCATTTGTCAACCAGGGTAGATG

Mouse Mir455-pair1/pair2-R: CTTAGGTACTCCCAGCATGGACGT

Expected PCR product size:

Mouse Mir455-pair1:

Wildtype allele: 538 bp

Mutant allele: ~490 bp, delete~40 bp

Mouse Mir455-pair2:

Wildtype allele: 538 bp

Mutant allele: ~490 bp, delete~40 bp

**3e. microRNA sequence**

Mouse Mir455 sequence: CTCCCTGGTGTGAGCGTATGTGCCTTTGGACTACATCGTGAACGCAGCACCATGCAGTCCACGGGCATATACACTTGCCTCA

*Note: The 5P and 3P mature sequences are colored in red.*

**3f. Off-target analysis**

*Note: Each gRNA has a quality score, with higher quality scores indicating greater specificity. Each potential off-target site also has a score, which is calculated based on the number of mismatches with the gRNA and the distances of the mismatches from PAM sequence, with high scores indicating greater potential that the off-target site will be recognized by the gRNA. The quality score of the gRNA is calculated based on the total number of off-target sites and their scores.*

Off-target analysis for gRNA1:

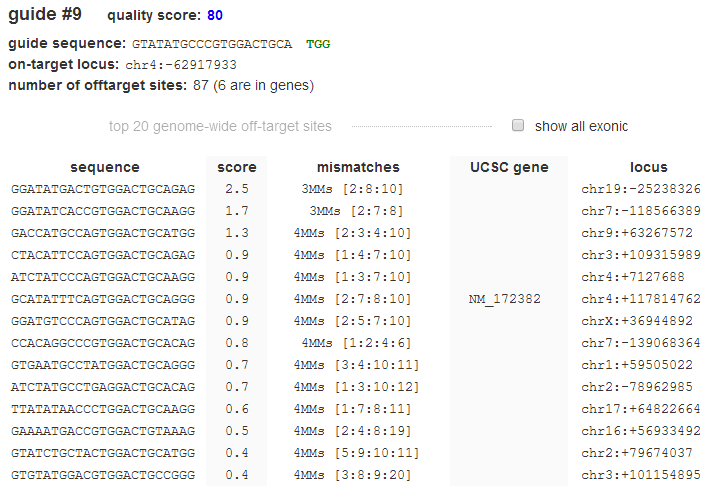

Off-target analysis for gRNA2:

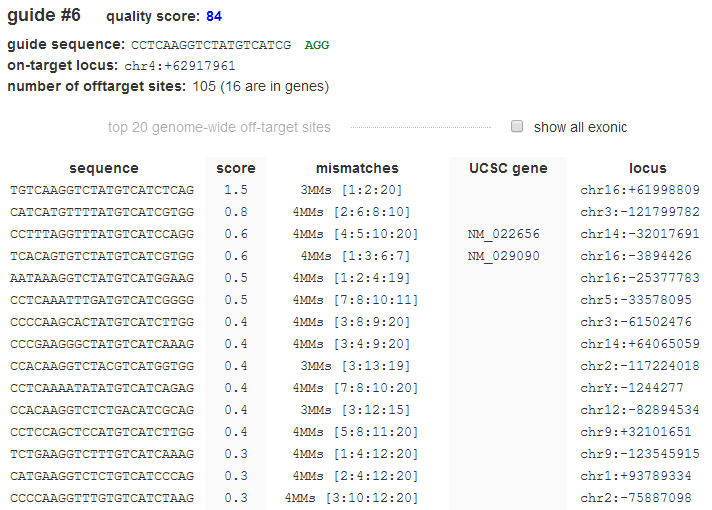

Off-target analysis for gRNA3:

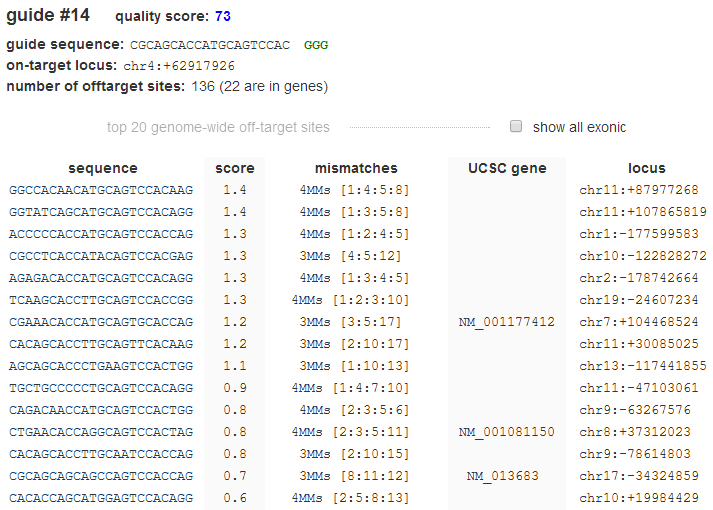

Off-target analysis for gRNA4:

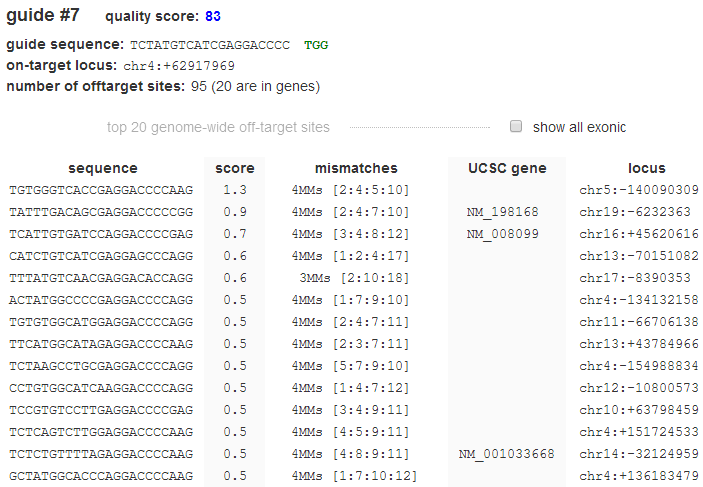
