## Supplemental Figure legends for "Novel microRNA-455-3p mouse models to study Alzheimer’s disease pathogenesis"

**Supplementary Figure Legends**

**SI Figure 1-** Schematic structure of miR-455-3p expression construct used to generate miR-455-3p transgenic mouse model.

**SI Figure 2-** Strategies for the deletion of miR-455-3p sequences from mouse genome and miR-455-3p knockout genotyping primers designing. Genomic regions of mouse miR-455 gene oriented from left to right; total size is 82 bp. The two pairs of genomic RNA construct showing binding site at 3’ prime of miR-455.

**SI Figure 3-** Sequence analysis of mouse miR-455 genomic regions in the miR-455-3p KO founders. In the mouse founder ID -10, the 41 bp, miR-455-3p genomic region was deleted and in mouse founder ID- 11, the 46 bp, miR-455-3p genomic region was deleted.

**SI Figure 4-** Confirmation of miR-455-3p TG pups by genotyping PCR showing specific product size of 297 bp and confirmation of miR-455-3p KO pups by genotyping PCR showing specific product size of 490 bp.

**SI Figure 5-** qRT-PCR analysis of key mitochondrial biogenesis genes (PGC1a, Nrf1, Nrf2 and TFAM), mitochondrial dynamics genes (Drp1, Fis1, OPA1, Mfn1 and Mfn2) and synaptic genes (SNAP25, PSD95 and MAP2) in WT controls (n=15), miR-455-3p TG (n=15) and miR-455-3p KO (n=15) mice.

**SI Figure 6-** Immunoblot analysis of key mitochondrial dynamics proteins in WT, TG and KO mice. Representative Immunoblots for Drp1, Fis1, Mfn1, Mfn2 and OPA1 proteins in WT (n=5), TG (n=5) and KO (n=5) mice.

**SI Figure 7-** Densitometry analysis for mitochondrial dynamics proteins (Drp1, Fis1, Mfn1, Mfn2 and OPA1) in WT, TG and KO mice. The Fis1 protein levels decreased significantly and Mfn1 and Opa1 proteins level was increased significantly in TG mice relative to WT mice

**SI Figure 8-** Immunoblot analysis of key synaptic proteins in WT, TG and KO mice. Representative Immunoblots for SNAP25, PSD95 and MAP2 proteins in WT (n=5), TG (n=5) and KO (n=5) mice.

**SI Figure 9-** Densitometry analysis for synaptic proteins (SNAP25, PSD95 and MAP2) in WT, TG and KO mice. All three synaptic proteins increased significantly with different significant levels in TG mice relative to WT and KO mice.

**SI Figure 10-** Immunostaining analysis of key mitochondrial dynamics proteins in WT (n=3), TG (n=3) and KO (n=3) mice. Representative Immunostaining images for Drp1, Fis1, OPA1, Mfn1 and Mfn2 proteins in WT, TG and KO mice. (10X magnification, 1mm scale).

**SI Figure 11-** Fluorescence intensity quantification for mitochondrial dynamics proteins (Drp1, Fis1, OPA1, Mfn1 and Mfn2) in WT, TG and KO mice.

**SI Figure 12-** Immunostaining analysis of key synaptic proteins in WT (n=3), TG (n=3) and KO (n=3) mice. Representative Immunostaining images for SNAP25, PSD95 and MAP2 proteins in WT, TG and KO mice.

**SI Figure 13-** Fluorescence intensity quantification for synaptic proteins (SNAP25, PSD95 and MAP2) in WT, TG and KO mice.

**SI Figure 14-** Immunostaining analysis of brain cells (Neuron, Astrocytes and Microglia) proteins in WT (n=3), TG (n=3) and KO (n=3) mice. Representative Immunostaining images for NeuN, GFAP, and Iba1 proteins in WT, TG and KO mice. (10X magnification, 500 µm scale).

**SI Figure 15-** Fluorescence intensity quantification for NeuN, GFAP, and Iba1 proteins in WT (n=3), TG (n=3) and KO (n=3) mice.

**SI Table 1-** Primers details used for qRT-PCR analysis

**SI Table 2-** Antibodies details used for immunoblotting and immunostaining analysis
