## Supplemental information for "Novel microRNA-455-3p mouse models to study Alzheimer’s disease pathogenesis"

**Morris Water Maze test**

The MWM apparatus consists of a 183 cm diameter pool filled with water to a depth of 48 cm. The pool temperature was maintained at 25 ± 0.5°C by the addition of warm water. The platform (18 cm diameter) was placed in the pool approximately 2 cm below the surface of the water in one of the four quadrants. On the four sides of the MWM apparatus, different shapes of visual cues were placed inside the wall of the pool in a diagonal pattern. The water in the pool was made opaque to obscure the platform location using non-toxic white tempera paint (Utrecht Art Supplies, Cranbury, NJ) (Vijayan et al., 2020). The pool was divided into four quadrants (compass locations: NE, NW, SW, and SE). The swimming animal was captured and recorded by using a video camera placed above the center of the pool, which is connected to a computer system running specialized tracking software (ANY-maze, Stoelting Co., USA). The platform was placed in the NE quadrant and remained at the same position during the whole experiment. Briefly, every group of animals was trained for four days in MWM, with four trials per day, with 15 minutes interatrial interval, so that one group of the animal was tested within 4 days/week. Each trial was a priest to run for one minute, but a trial ended once the animal was positioned on the platform for 3 seconds. If the animal didn’t find the platform within one minute, they were placed on the platform using the net for 3 seconds. After every trial animal was dried with a towel and placed into a holding cage. The day after four days of training, the platform was removed, and one probe trial was conducted to evaluate the time spent in the previously correct quadrant platform. Two hours later, the reversal trial was performed. The platform was placed opposite to the previous location. The animal was tested to run one minute, but a trial ended once the animal was positioned on the platform for 3 seconds. Time to find the platform and percentage of time spent in each quadrant were determined from the retrieved videotapes. All animals were trained 4 times, later assessed for another 6 times for latency to find the platform.
